# Multi-region exome sequencing of ovarian teratomas reveals 2N near-diploid genomes, paucity of somatic mutations, and extensive allelic imbalances shared across mature, immature, and disseminated components

**DOI:** 10.1101/818534

**Authors:** Michael B. Heskett, John Z. Sanborn, Christopher Boniface, Benjamin Goode, Jocelyn Chapman, Karuna Garg, Joseph T. Rabban, Charles Zaloudek, Stephen C. Benz, Paul T. Spellman, David A. Solomon, Raymond J. Cho

## Abstract

Immature teratoma is a subtype of malignant germ cell tumor of the ovary that occurs most commonly in the first three decades of life, frequently with bilateral ovarian disease. Despite being the second most common malignant germ cell tumor of the ovary, little is known about its genetic underpinnings. Here we performed multi-region whole exome sequencing to interrogate the genetic zygosity, clonal relationship, DNA copy number, and mutational status of 52 pathologically distinct tumor components from 10 females with ovarian immature teratomas, with bilateral tumors present in 5 cases and peritoneal dissemination in 7 cases. We found that ovarian immature teratomas are genetically characterized by 2N near-diploid genomes with extensive loss of heterozygosity and an absence of genes harboring recurrent somatic mutations or known oncogenic variants. All components within a single ovarian tumor (immature teratoma, mature teratoma with different histologic patterns of differentiation, and yolk sac tumor) were found to harbor an identical pattern of loss of heterozygosity across the genome, indicating a shared clonal origin. In contrast, the 4 analyzed bilateral teratomas showed distinct patterns of zygosity changes in the right versus left sided tumors, indicating independent clonal origins. All disseminated teratoma components within the peritoneum (including gliomatosis peritonei) shared a clonal pattern of loss of heterozygosity with either the right or left primary ovarian tumor. The observed genomic loss of heterozygosity patterns indicate that diverse meiotic errors contribute to the formation of ovarian immature teratomas, with 11 out of the 15 genetically distinct clones determined to result from the failure of meiosis I or II. Overall, these findings suggest that copy-neutral loss of heterozygosity resulting from meiotic abnormalities may be sufficient to generate ovarian immature teratomas from germ cells.

## Introduction

Germ cell tumors (GCTs) are a diverse group of neoplasms that display remarkable heterogeneity in their anatomical site, histopathology, prognosis, and molecular characteristics [1]. GCTs can occur in the ovaries, testes, and extragonadal sites, with the most common extragonadal locations being the anterior mediastinum, retroperitoneum, and intracranially in the pineal region [2]. GCTs are classified by the World Health Organization into seven histological subtypes: mature teratoma, immature teratoma, seminoma/dysgerminoma/germinoma (depending on site of origin in the testis, ovary, or extragonadal), yolk sac tumor, embryonal carcinoma, choriocarcinoma, and mixed germ cell tumor [3].

GCTs are the most common non-epithelial tumors of the ovary, but only account for approximately 3% of all ovarian cancers [4]. Approximately 90% of ovarian GCTs are composed entirely of mature teratoma (commonly termed “dermoid cyst”), which is the only benign subtype of ovarian GCT. Among the malignant subtypes, dysgerminoma is the most common and immature teratoma is the second most common. Ovarian teratomas contain tissue elements from at least 2 of the 3 germ cell layers and frequently display a disorganized mixture of mature tissues including skin and hair (ectoderm), neural tissue (ectoderm), fat (mesoderm), muscle (mesoderm), cartilage (mesoderm), bone (mesoderm), respiratory epithelium (endoderm), and gastrointestinal epithelium (endoderm). Teratomas can occur in the mature form, composed exclusively of mature tissues, or the immature form, which contains variable amounts of immature elements (usually primitive neuroectodermal tissue consisting of primitive neural tubules) in a background of mature teratoma [5]. Not infrequently, malignant GCTs of the ovary contain a mixture of different histologic subtypes (*e.g.* both dysgerminoma and yolk sac tumor), for which the designation mixed germ cell tumor is used, often with the approximate fraction of each histologic subtype specified by the diagnostic pathologist. Extensive tissue sampling and microscopic review of ovarian GCTs are required to appropriately evaluate for the presence of admixed malignant subtypes, which is critical for appropriately guiding prognosis and patient management.

The majority of ovarian GCTs (57%) are confined to the ovary at time of diagnosis (stage I) which confers a 99% 5-year survival [4]. Even when distant metastases are present at time of diagnosis (stage IV), 5-year survival of ovarian GCT is relatively high at 69% [4]. This long-term survival in females even with disseminated or metastatic ovarian GCTs reflects the sensitivity of these tumors to the standard cytotoxic chemotherapy regimen of bleomycin, etoposide, and cisplatin [1].

Somatic mutation and DNA copy number analysis of testicular GCTs has now been performed by The Cancer Genome Atlas Research Network and several other groups [6-14]. These analyses have revealed a very low mutation rate (approximately 0.3 somatic mutations per Mb) and only three genes harboring recurrent somatic mutations at significant frequency (*KIT, KRAS*, and *NRAS*), in which mutations are exclusively present in seminomas but not non-seminomatous GCTs [7-14]. Copy number analysis has revealed that testicular GCTs are often hyperdiploid, with the majority (>80%) harboring isochromosome 12p or polysomy 12p that is present in both seminomas and non-seminomatous GCTs [6, 9, 12, 13, 14]. Similar oncogenic *KIT* and *KRAS* mutations as well as polysomy 12p have also been frequently found in ovarian dysgerminomas and intracranial germinomas, indicating a shared molecular pathogenesis with testicular seminomas [15-20].

Beyond dysgerminomas, few studies have performed genome-level analysis of ovarian GCTs, and the genetic basis of ovarian teratomas (both mature and immature forms) remains unknown. Polysomy 12 and *KIT* mutations have been found in ovarian mixed germ cell tumors containing a dysgerminoma component, but have not been identified in pure teratomas [20]. Early studies of ovarian mature teratomas reported that tumor karyotypes were nearly always normal (*i.e.* 46,XX), but chromosomal zygosity markers were often homozygous in the tumor [21-24]. This loss of heterozygosity may be explained by the hypothesis that teratomas and other germ cell tumors arise from primordial germ cells due to one of five different plausible meiotic abnormalities, each producing distinct chromosomal patterns of homozygosity [23-26]. Parthenogenesis (from the Greek *parthenos*: ‘virgin’, and *genesis*: ‘creation’) is used to describe the development of germ cell tumors from unfertilized germ cells via these different mechanisms of origin, which potentially include failure of meiosis I, failure of meiosis II, whole genome duplication of a mature ovum, and fusion of two ova. However, no studies to date have used genome-level sequencing analysis to identify the specific parthenogenetic mechanism giving rise to individual ovarian GCTs.

Here we present the results of multi-region whole exome sequencing of 52 pathologically distinct tumor components from 10 females with ovarian immature teratomas, with bilateral tumors present in 5 cases and peritoneal dissemination in 7 cases. Our analyses define ovarian immature teratoma as a genetically distinct entity amongst the broad spectrum of human cancer types studied to date, which is characterized by a 2N near-diploid genome, paucity of somatic mutations, and extensive allelic imbalances. Our results further shed light on the parthenogenetic origin of ovarian teratomas and reveal that diverse meiotic errors are likely to drive development of this germ cell tumor.

## Materials and Methods

### Study population and tumor specimens

This study was approved by the Institutional Review Board of the University of California, San Francisco. Ten patients who underwent resection of ovarian immature teratomas at the University of California, San Francisco Medical Center between the years 2002-2015 were included in this study. All tumor specimens were fixed in 10% neutral-buffered formalin and embedded in paraffin. Pathologic review of all tumor specimens was performed to confirm the diagnosis by a group of expert gynecologic pathologists (K.G., J.T.R., C.Z., and D.A.S.).

### Whole exome sequencing

Tumor tissue from each of the indicated ovarian and disseminated germ cell tumor components was selectively punched from formalin-fixed, paraffin-embedded blocks using 2.0 mm disposable biopsy punches (Integra Miltex Instruments, cat# 33-31-P/25). These punches were made into areas histologically visualized to be composed entirely of the indicated germ cell component (*e.g.* immature teratoma, mature teratoma, yolk sac tumor, gliomatosis peritonei). Uninvolved normal fallopian tube was also selectively punched from formalin-fixed, paraffin-embedded blocks as a source of constitutional DNA for each of the ten patients. Genomic DNA was extracted from these tumor and matched normal tissue samples using the QIAamp DNA FFPE Tissue Kit (Qiagen) according to the manufacturer’s protocol. 500 ng of genomic DNA was used as input for capture employing the xGen Exome Research Panel v1.0 (Integrated DNA Technologies). Hybrid-capture libraries were sequenced on an Illumina HiSeq 4000 instrument.

### Mutation calling and loss of heterozygosity analysis

Sequence reads were aligned to the hg19 reference genome using Burrows-Wheeler Alignment tool [27]. Duplicate reads were removed and base quality scores recalibrated with GATK prior to downstream analysis [28]. Candidate somatic mutations were identified with MuTect v1.1.5 with the minimum mapping quality parameter set to 20. dbSNP build 150 was used to identify and remove SNPs. The following additional filters were applied to candidate mutations from MuTect output: minimum tumor depth 30, minimum normal depth 15, minimum variant allele frequency 15%, maximum variant allele presence in normal 2%. Finally, all candidate mutations were manually reviewed in the Integrative Genome Viewer to remove spurious variant calls likely arising from sequencing artifact [29, 30]. FACETS was used to determine allele-specific copy number and loss of heterozygosity regions across the genome [31]. To determine genetic mechanism of origin, tumors were classified into one of five plausible categories based on the zygosity status at centromeric and distal regions, as described by Surti et al [25]. For visualization of zygosity changes across the genome in the tumor specimens, the absolute difference between theoretical heterozygosity (allele frequency = 0.5) of tumor versus normal was plotted.

## Results

### Patient cohort

Clinical data from the patient cohort is summarized in Table 1. The 10 females ranged in age at time of initial surgery from 8-29 years (median 17 years). None were known to have Turner syndrome or other gonadal dysgenesis disorder, nor any known familial tumor predisposition syndrome. All patients underwent resection of a primary ovarian mass, along with debulking of disseminated disease observed in the peritoneum at time of initial oophorectomy for 5 patients. Bilateral ovarian tumors were present in 5 of the 10 patients, 2 with synchronous disease at time of initial diagnosis (a and b) and 3 with metachronous disease that was identified and resected during the period of clinical follow-up (g, h, and j). Primary ovarian tumor size ranged from 4-30 cm (median 15 cm). Four of the patients were treated with adjuvant chemotherapy using bleomycin, etoposide, and cisplatin after initial surgery based on the presence of disseminated immature teratoma in the peritoneum (a, b, e, and k). A fifth patient was treated with adjuvant chemotherapy using bleomycin, etoposide, and cisplatin following resection of a synchronous ovarian immature teratoma at 4.8 years after resection of a contralateral ovarian mature teratoma (g). One exceptional 14-year-old patient (d) initially underwent resection of a unilateral 18 cm ovarian immature teratoma and debulking of disseminated peritoneal disease. Subsequent PET/CT showed widespread bulky lymphadenopathy. She underwent resection of a supraclavicular lymph node at 0.6 years after initial oophorectomy which contained metastatic primitive neuroectodermal tumor (PNET) and atypical gliomatosis histologically resembling an anaplastic astrocytoma of the central nervous system. She was treated with intensive multiagent chemotherapy including vincristine, doxorubicin, cyclophosphamide, ifosfamide, and etoposide. Over the next three years, she underwent additional resections of recurrent/progressive disease in the peritoneum, cyberknife radiotherapy to left axilla, and multiple courses of chemotherapy, first with temozolomide and then with cyclophosphamide and topotecan. She remains alive with stable disease at last clinical follow-up (6.6 years after initial surgery). All other patients in this cohort also remain alive with stable disease or without evidence of disease recurrence at last clinical follow-up (range 2.4-15.3 years, median 6.6 years, excluding patient i with no clinical follow-up data after initial resection).

### Histologic features of the ovarian immature teratomas

Pathologic diagnosis for the ovarian germ cell tumors is summarized in Table 1, and representative photomicrographs are shown in Figure 1. All 10 patients had primary ovarian immature teratomas composed of primitive neural tubules in a background of mature teratoma. In 2 patients, there were additionally admixed small foci of yolk sac tumor and embryonal carcinoma, thereby warranting designation as mixed germ cell tumor, although mature and immature teratoma were the predominant elements in both cases. Five patients also had teratomas involving the contralateral ovary, 2 of which were synchronous and 3 of which were metachronous. The contralateral ovarian tumors were also immature teratomas in 2 patients (a and h), whereas the contralateral ovarian tumors were composed entirely of mature teratoma in 3 patients (b, g, and j). Disseminated disease was found in the peritoneum of 7 patients, which consisted of a combination of immature and mature elements in 5 patients and mature elements only in 2 patients. The disseminated immature elements in one of these patients (d) was histologically diagnosed as primitive neuroectodermal tumor (PNET), as it was composed of sheets of primitive small round blue cells with diffuse immunoreactivity for synaptophysin and without organization into neural tubules or evidence of neuroglial differentiation. Six patients had peritoneal implants composed of mature glial tissue that has been termed gliomatosis peritonei. This gliomatosis peritonei was of low cellularity and composed of cytologically bland glial cells in 5 patients, whereas the gliomatosis peritonei was hypercellular and composed of cytologically atypical glial cells resembling anaplastic astrocytoma of the central nervous system in 1 patient (d).

**Fig. 1.**
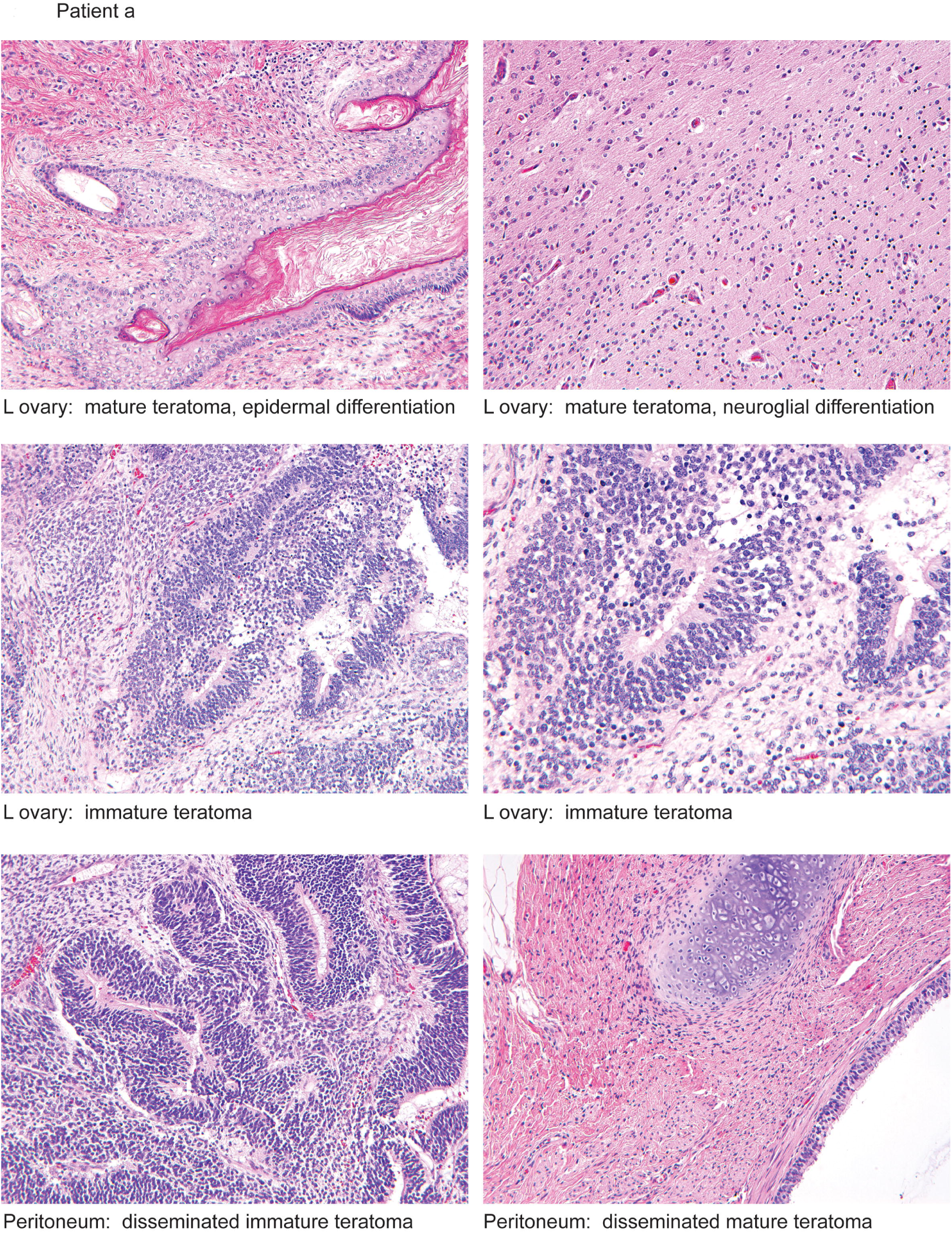
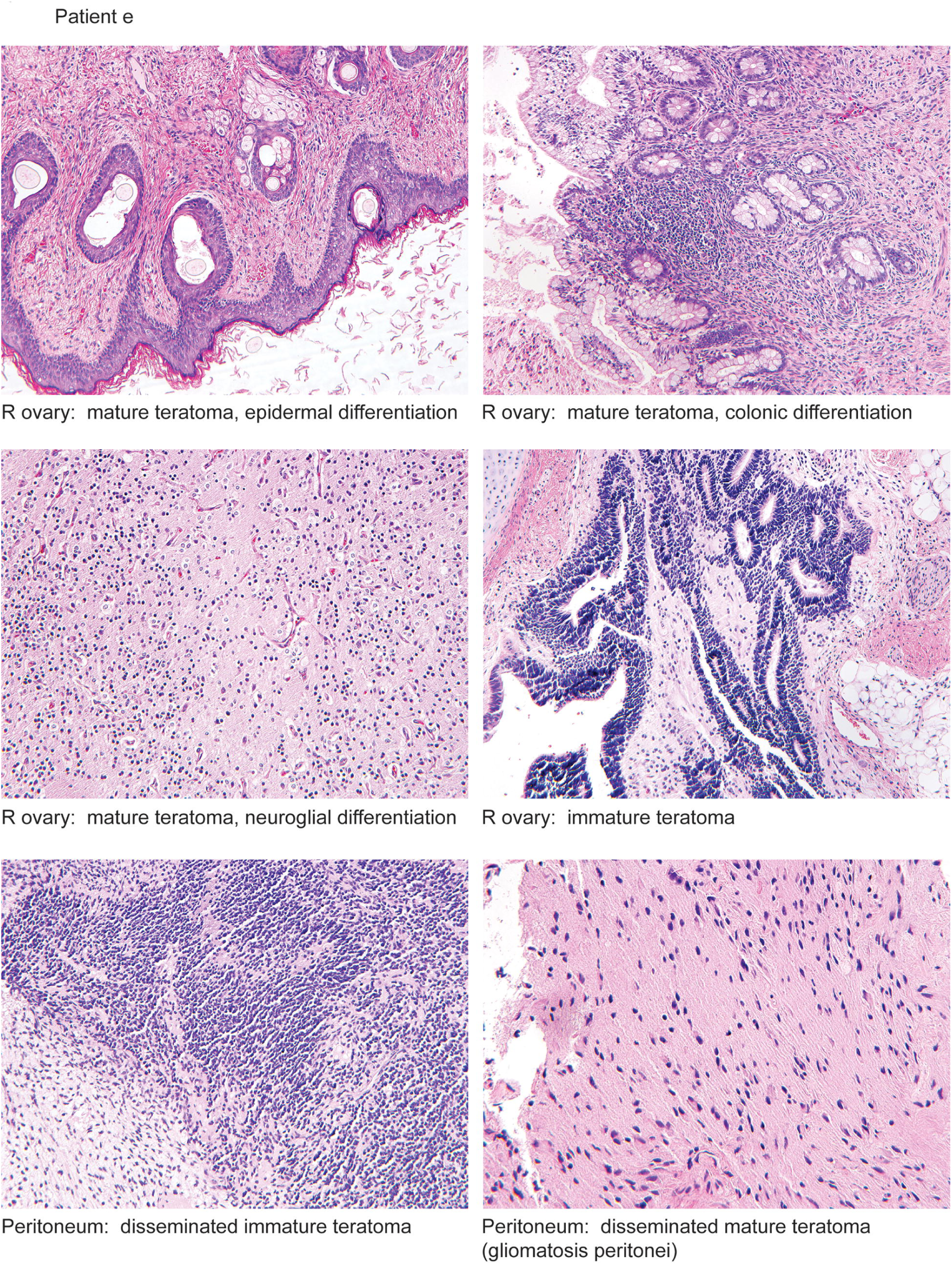
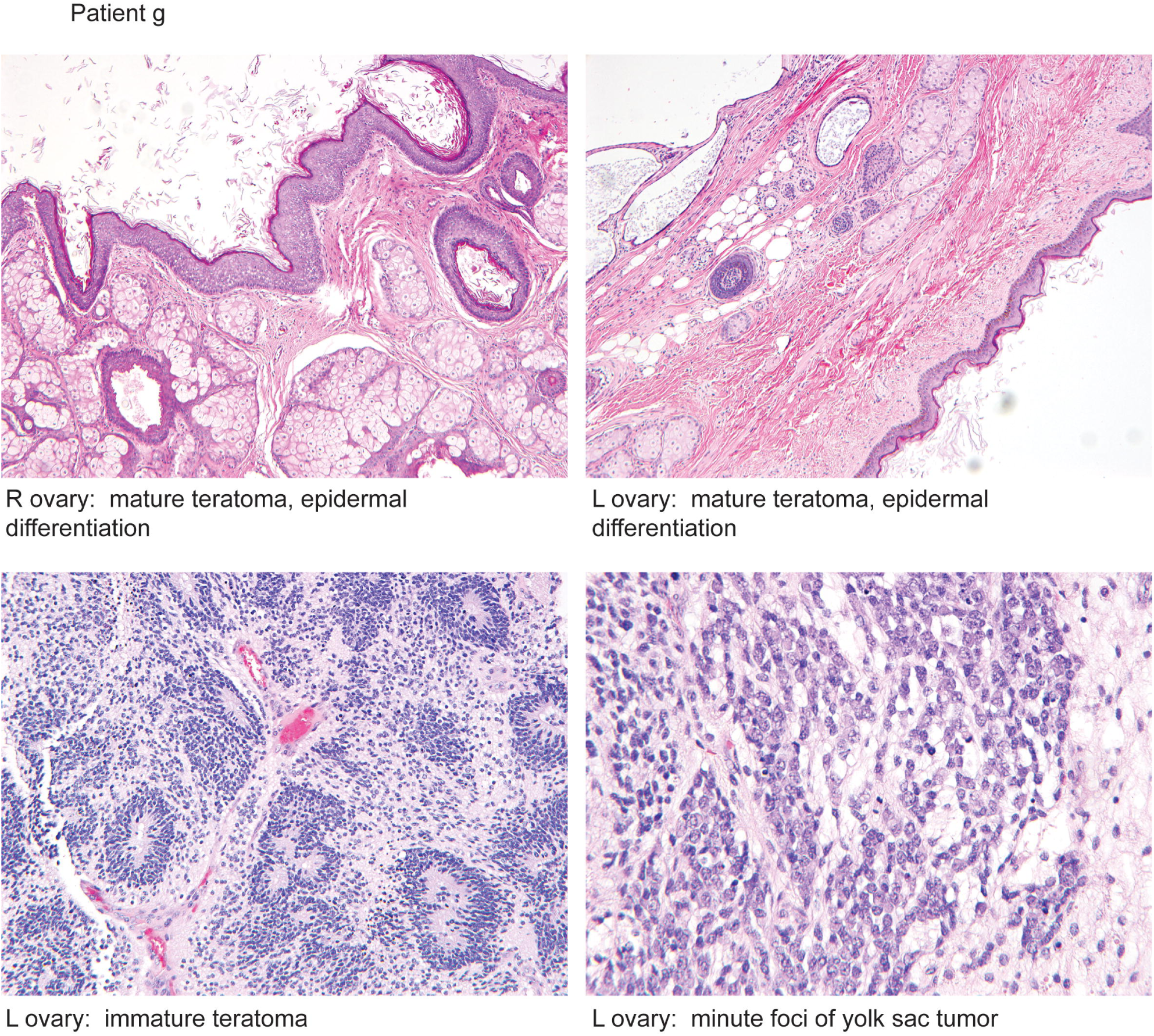
Histology images of the ovarian immature teratomas from three representative patients that were studied by whole exome sequencing. Shown are hematoxylin and eosin stained sections illustrating the different tumor regions from the primary ovarian mass as well as disseminated disease in the peritoneum from which genomic DNA was selectively extracted for analysis. **a** Patient a is a 16-year-old female who underwent resection of synchronous bilateral ovarian immature teratomas and debulking of disseminated peritoneal disease. **b** Patient e is a 29-year-old female who underwent resection of a unilateral ovarian immature teratoma and debulking of disseminated peritoneal disease. **c** Patient g is a 25-year-old female who underwent resection of a unilateral ovarian immature teratoma and then four years later underwent resection of a contralateral ovarian mature teratoma (no immature component present).

### Multi-region whole exome sequencing of ovarian immature teratomas

Genomic DNA was extracted from 52 tumor regions consisting of ovarian immature teratoma, mature teratoma, yolk sac tumor, and disseminated teratomatous elements, along with uninvolved normal fallopian tube tissue from the 10 female patients (Table 2). Hybrid exome capture and massively parallel sequencing by synthesis on an Illumina platform was performed to an average depth of 203x per sample, as described in the Methods. Sequencing metrics are displayed in Supplementary Table 1. The number of tumor regions sequenced per patient ranged from 2 to 9, with a median of 4.

### Paucity of somatic single nucleotide variants in ovarian immature teratomas

Based on this whole exome sequencing of 52 tumor samples, we identified a total of only 31 unique high-confidence somatic nonsynonymous mutations (Supplementary Table 2). Despite high sequencing depth, we detected somatic nonsynonymous mutations in only 21 of the 52 samples, and the average number of somatic nonsynonymous mutations in the mutated samples was 0.8 per exome. The mean somatic mutation burden (commonly also referred to as total mutation burden or TMB) per tumor sample was 0.02 non-synonymous mutations per Mb, which is among the lowest of any human cancer type that has been analyzed to date.

Only 1 of the somatic nonsynonymous mutations (*PKFP* p.A158V in patient a, RefSeq transcript NM_002627) was present in all tumor regions sequenced from a single patient, thereby indicating its clonality and acquisition early during tumorigenesis. However, the other 30 somatic nonsynonymous mutations were present only in a single tumor region or a subset of the tumor regions sequenced, thereby indicating their subclonality and acquisition later during tumorigenesis. For example, the *PKFP* p.A158V mutation was present in all tumor regions sequenced from patient a, including the immature and both mature teratoma components from the left ovary, as well as the disseminated immature teratoma, mature teratoma, and yolk sac tumor components in the peritoneum. In contrast, the *CCS* p.R112C (RefSeq transcript NM_005125) mutation was exclusively present in the ovarian mature teratoma component with neuroglial differentiation, and the *CIITA* p.R2C (RefSeq transcript NM_000246) mutation was only present in the disseminated immature teratoma and yolk sac tumor components. Thus, none of these 30 somatic nonsynonymous mutations could have plausibly been the initiating genetic driver in this cohort of ovarian immature teratomas.

No genes were identified to harbor recurrent somatic nonsynonymous mutations across the 10 patients (*i.e.* no gene was mutated in more than a single patient). Furthermore, no well-described oncogenic variants (e.g. *BRAF* p.V600E) were identified in any of the 52 tumors samples. Among the 723 genes currently annotated in the Cancer Gene Census of the Catalog of Somatic Mutations in Cancer (COSMIC) database version 90 release, only 4 were identified to harbor somatic nonsynonymous mutations in this ovarian immature teratoma cohort. However, the variants in these 4 genes (*TP53, NF1, CTNNB1*, and *NOTCH2*) were each found in a single tumor sample in this cohort, were all non-truncating missense variants, and are not known recurrent somatic mutations in the current version of the COSMIC database. Thus, the functional significance of the identified mutations in these 4 genes is uncertain, and they may likely represent bystander alterations rather than driver mutations. Although *KIT, KRAS, NRAS*, and *RRAS2* are recurrently mutated oncogenes that drive ovarian dysgerminomas and testicular germ cell tumors [14, 19, 20], we found no mutations in these genes in this cohort of ovarian immature teratomas.

### Ovarian immature teratomas have 2N diploid or near-diploid genomes with extensive loss of heterozygosity

Using FACETS to infer copy number status and the genotype data of common polymorphisms from the exome sequencing, we next assessed the chromosomal copy number and zygosity status of the 52 tumor samples (Table 2). All of the 52 tumor samples were found to harbor 2N diploid or near-diploid genomes. All tumor samples from 6 of the patients had normal 46,XX diploid genomes. All tumor samples from 3 of patients had near-diploid genomes with clonal gain of a single whole chromosome (+3 in patient d, +14 in patient i, and +10 in patient k). In patient b with bilateral ovarian teratomas, the mature teratoma from the right ovary harbored a normal 46,XX diploid genome, whereas all tumor samples from the left ovary and all disseminated peritoneal tumor samples harbored near-diploid genomes with clonal gains of whole chromosomes 3 and X. No focal amplification or deletion events were identified in any of the 52 tumor samples. None of the tumor samples harbored isochromosome 12p or polysomy 12p.

We next plotted the absolute change in allele frequency (*ΔAF)* for the 52 tumor samples based on the genotype of common polymorphisms across each of the chromosomes, using an average of approximately 17,000 informative loci per genome. Whereas an allele frequency of 0.5 equals the normal heterozygous state for a diploid genome, an allele frequency of 0.0 or 1.0 equals a homozygous state, which could be due to either chromosomal copy loss or copy-neutral loss of heterozygosity. We observed extensive copy-neutral loss of heterozygosity across the genomes of each of the 52 tumor samples from all 10 patients (Figure 2).

**Fig. 2.**
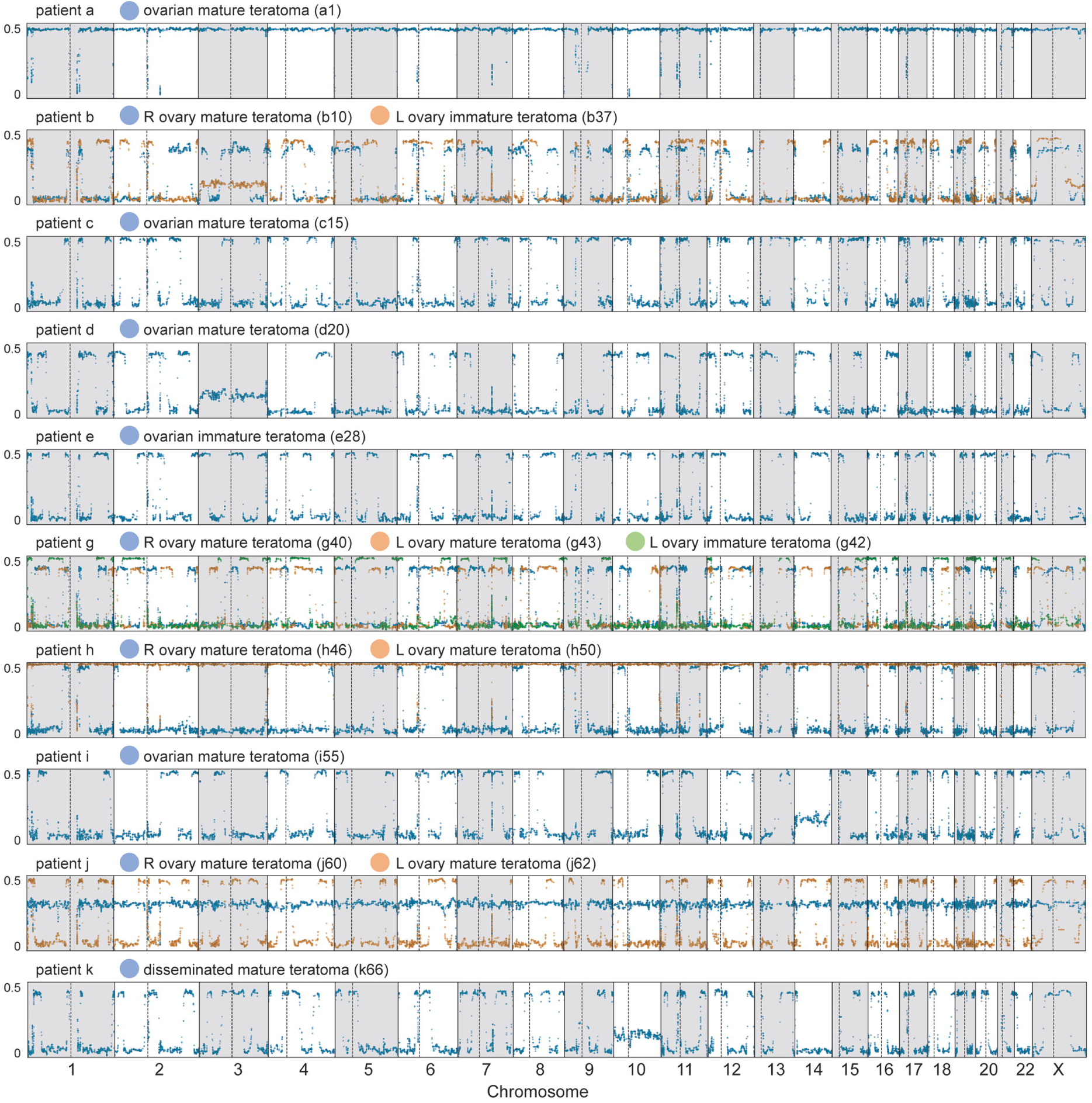
Ovarian immature teratomas are characterized by extensive genomic loss of heterozygosity. Plots of Δallele frequency (ΔAF) were generated from the whole exome sequencing data for each of the 52 tumor regions from 10 patients with ovarian immature teratomas. Two distinct tumor clones were identified in four patients (b, g, h, and j) who all had bilateral ovarian teratomas. While all tumor regions harbored a 2N diploid or near-diploid genome, extensive genomic loss of heterozygosity was observed in each of the different tumor components analyzed. Each point represents one informative polymorphic locus. Points near the top of the y-axis represent single nucleotide polymorphisms that are homozygous in the tumor, whereas points near the bottom of the y-axis are heterozygous. y-axis, ΔAF. x-axis, chromosome. Dotted line, centromere. ΔAF is calculated as the absolute difference between theoretical heterozygosity (AF=0.5).

### Identical patterns of genomic loss of heterozygosity among mature, immature, and disseminated components in an ovarian teratoma confirm a single clonal origin

We next compared the regions of the genome affected by copy-neutral loss of heterozygosity among the different tumor regions sequenced for each individual patient. In the 5 females with unilateral ovarian disease (patients c, d, e, i, and k), we observed the identical pattern of allelic imbalance across the genome in each of the different tumor components, including immature teratoma, mature teratoma with different histologic patterns of differentiation, and disseminated teratomatous elements in the peritoneum. These results confirm a single clonal origin for all teratomatous components, both in the primary ovarian tumor and disseminated in the peritoneum, for women with unilateral ovarian immature teratomas.

### Bilateral ovarian teratomas originate independently

Four patients in this cohort (b, g, h, and j) had bilateral ovarian teratomas that were both independently sequenced and analyzed for patterns of copy-neutral loss of heterozygosity across the genome. We found that tumors from the left and right ovaries had different patterns of allelic imbalance across the genome in each of the different tumor components studied, providing evidence that bilateral ovarian teratomas originate independently. Furthermore, all of the peritoneal disseminated components harbored a pattern of allelic imbalance that was identical to one of the two ovarian tumors, enabling deduction of the specific ovarian tumor from which the disseminated disease was clonally related. For example, patient h is an 8-year-old girl who initially underwent resection of a 17 cm immature teratoma from the left ovary, and then 9 years later underwent resection of a 16 cm immature teratoma from the right ovary as well as debulking of disseminated disease in the peritoneum (gliomatosis peritonei). The immature teratoma and two mature teratoma regions studied from the left ovary had an identical pattern of allelic imbalance, whereas the immature teratoma and two mature teratoma regions studied from the right ovary had an identical pattern of allelic imbalance that was distinct from the tumor elements in the contralateral ovary. Additionally, the gliomatosis peritonei had an identical pattern of allelic imbalance as the immature and mature teratoma components from the right ovary (Figure 3).

**Fig. 3.**
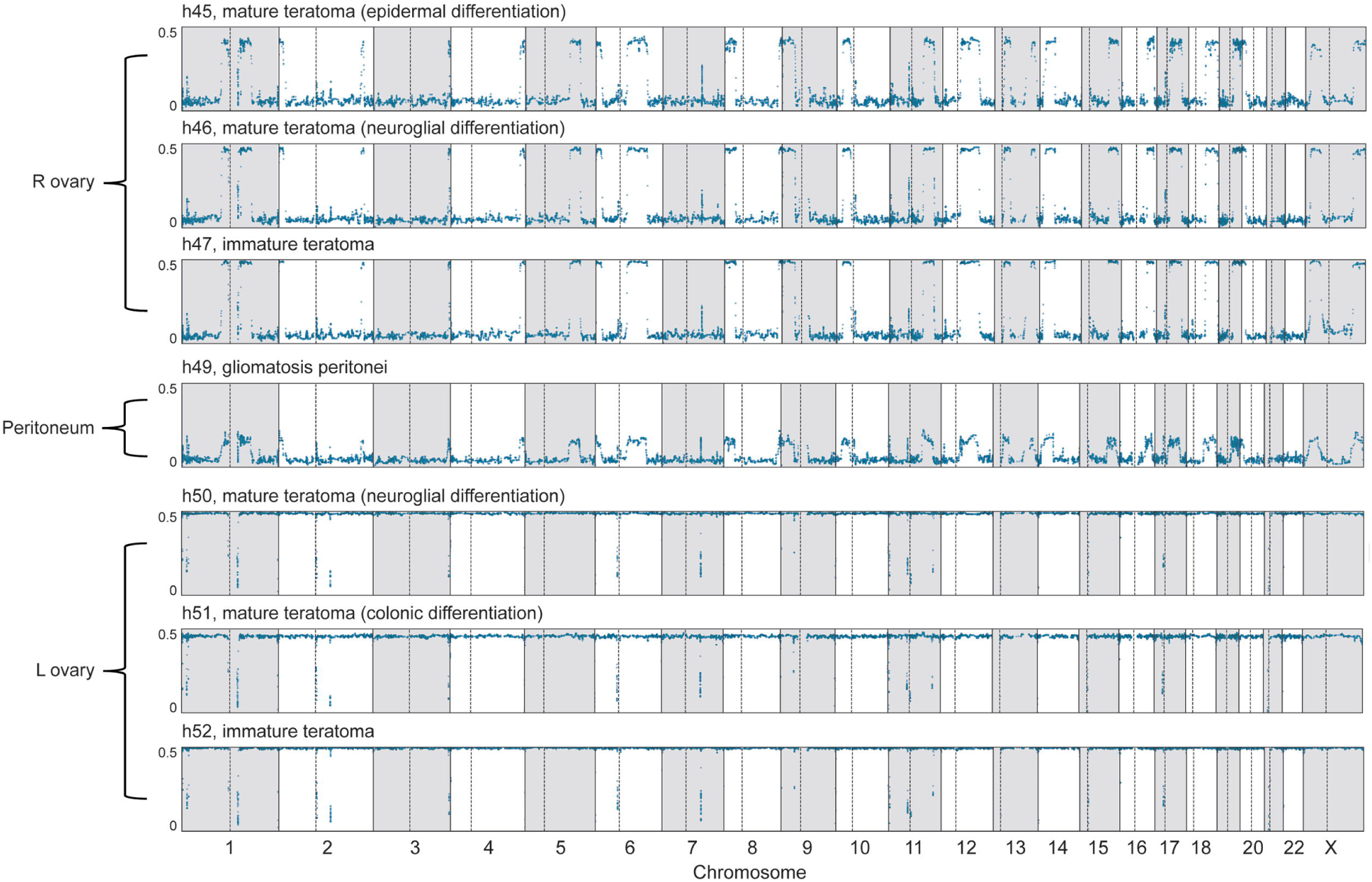
Identical patterns of genomic loss of heterozygosity among all mature, immature, and disseminated components in ovarian teratomas confirm a single clonal origin, except in females with bilateral tumors. Plots of Δallele frequency (ΔAF) were generated from the whole exome sequencing data for each of the 7 different tumor regions from patient h, an 8-year-old girl who initially underwent resection of a 17 cm immature teratoma from the left ovary, and then 9 years later underwent resection of a 16 cm immature teratoma from the right ovary as well as debulking of disseminated disease in the peritoneum (gliomatosis peritonei). While all tumor regions harbored a diploid genome, extensive genomic loss of heterozygosity was observed in each of the different tumor components. The immature teratoma and two mature teratoma regions studied from the left ovary had the identical pattern of allelic imbalance, whereas the immature teratoma and two mature teratoma regions studied from the right ovary shared an identical pattern of allelic imbalance that was distinct from the tumor elements in the contralateral ovary. Additionally, the gliomatosis peritonei had an identical pattern of allelic imbalance as the immature and mature teratoma components from the right ovary. Each point represents one informative polymorphic locus. Points near the top of the y-axis represent single nucleotide polymorphisms that are homozygous in the tumor, whereas points near the bottom of the y-axis are heterozygous. y-axis, ΔAF. x-axis, chromosome. Dotted line, centromere. ΔAF is calculated as the absolute difference between theoretical heterozygosity (AF=0.5).

### Patterns of genomic loss of heterozygosity in ovarian immature teratomas can be used to deduce meiotic error mechanism of origin

Five distinct parthenogenetic mechanisms of origin have been proposed to describe the development of germ cell tumors from unfertilized germ cells, which include failure of meiosis I, failure of meiosis II, whole genome duplication of a mature ovum, and fusion of two ova. Distinct chromosomal zygosity patterns are predicted to result from each of these different mechanisms [25], which are illustrated in Figure 4. We used the chromosomal zygosity patterns from the whole exome sequencing data to deduce the meiotic mechanism of origin for the 15 distinct tumor clones identified in the 10 female patients. Five of the tumor clones were deduced to result from failure of meiosis I, 6 from failure of meiosis II, 3 from whole genome duplication of a mature ovum, and 1 from fusion of two ova (Table 2). These findings indicate that failure at multiple stages during germ cell development can contribute to the development of ovarian teratomas.

**Fig. 4.**
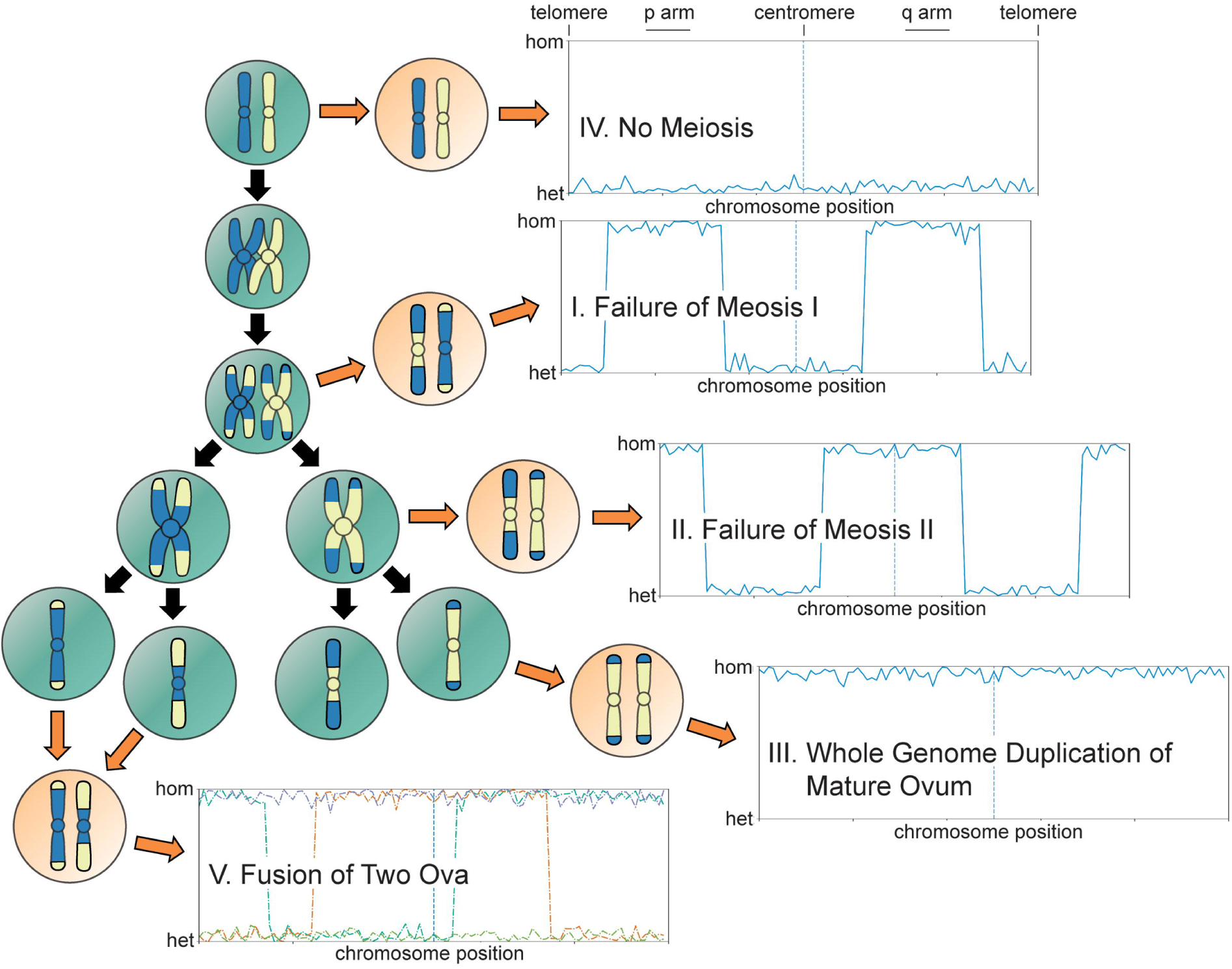
The five proposed genetic mechanisms of origin of ovarian teratomas from a germ cell. One homologous chromosome pair undergoing two genetic crossing over events is illustrated for simplicity. Orange arrows depict aberrant outcomes of meiosis. Black arrows depict the normal path through meiosis. Each plot depicts a simulated example of the chromosomal loss-of-heterozygosity pattern that arises from each of the five hypothetical mechanisms of origin, measured by the allele frequency difference of SNPs in the tumor compared to constitutional DNA. Y-axis: the two possible zygosity states in a diploid cell (top = homozygosity, bottom = heterozygosity). X-axis: position along an individual chromosome. Vertical dotted blue line depicts the centromere. For Mechanism V, the homozygosity pattern on each chromosome will vary based on the number and location of crossing over events. Adapted from Surti et al. [25].

## Discussion

We present the first multi-region exome sequencing analysis of ovarian immature teratomas including mature, immature, and disseminated components. We report a strikingly low abundance of somatic mutations and infrequent copy number aberrations, without pathogenic mutations identified in any well-described oncogenes or tumor suppressor genes, as well as an absence of any novel genes harboring recurrent mutations across the cohort. We generated high-resolution zygosity maps of ovarian teratomas that deepen understanding of the parthenogenetic mechanisms of origin of ovarian teratomas from primordial germ cells originally proposed nearly 50 years ago [21]. Ovarian teratoma is genetically unique among all human tumor types studied to date given its extremely low mutation rate and extensive genomic loss of heterozygosity. Our findings suggest that meiotic non-disjunction events producing a 2N near-diploid genome with extensive allelic imbalances are responsible for the development of ovarian immature teratomas.

Analysis of the multi-region exome sequencing data was used to study the clonal relationship of immature and mature teratoma elements, as well as admixed foci of yolk sac tumor, and also disseminated teratoma in the peritoneum. We find that all these different tumor components are indistinguishable based on chromosomal copy number alterations and loss of heterozygosity patterns, indicating a shared clonal origin. This finding suggests that epigenetic differences are likely responsible for the striking variation in differentiation patterns in teratomas.

Notably, gliomatosis peritonei is a rare phenomenon in which deposits of mature glial tissue are found in the peritoneum, which occurs almost exclusively in association with immature teratoma of the gonads [32, 33]. Two theories currently exist to explain the origin of gliomatosis peritonei: the first being that it is derived from peritoneal dissemination of teratoma with differentiation into mature glial cells, and the other being spontaneous metaplasia of peritoneal stem cells to glial tissue in individuals with gonadal teratoma [34, 35]. A prior study of five samples had concluded that gliomatosis peritonei was genetically unrelated to the primary ovarian teratoma based on zygosity analysis of a small number of microsatellite markers [35]. However, our study based on genotyping data from thousands of informative polymorphic loci unequivocally demonstrates that gliomatosis peritonei is clonally related to the ovarian primary immature teratoma in all cases, thereby confirming the first theory of origin.

While all ovarian and disseminated tumor components in the 5 patients with unilateral disease in this cohort were found to be clonally related, 4 patients had bilateral ovarian teratomas that were independently analyzed and found to have distinct clonal origins. We found that tumors from the left and right ovaries had different patterns of loss of heterozygosity across the genome in each of the different tumor components that were sequenced, providing evidence that bilateral ovarian teratomas originate independently. Additionally, all of the disseminated components in the peritoneum harbored a pattern of allelic imbalance that was identical to one of the two ovarian tumors, enabling assignment of origin to the specific ovarian primary tumor. Why a significant proportion of women with ovarian teratomas also develop genetically independent teratomas in the contralateral ovary (either synchronously or metachronously) remains undefined. Analysis of the constitutional DNA sequence data from the 10 patients in our cohort, 5 of whom had bilateral ovarian teratomas, did not identify pathogenic variants in the germline known to be associated with increased cancer risk. However, the possibility of an unidentified germline risk allele(s) responsible for teratoma development remains a possibility. Given the extensive loss of heterozygosity across the genomes of ovarian teratomas, pinpointing any single responsible gene amongst the numerous common regions of allelic imbalance is a significant obstacle.

In summary, our multi-region whole exome sequencing analysis of ovarian immature teratomas has revealed that multiple different meiotic errors can give rise to these genetically distinct tumors that are characterized by extensive allelic imbalances and a paucity of somatic mutations and copy number alterations.

## Acknowledgements

This study was supported by NIH Director’s Early Independence Award (DP5 OD021403) and the UCSF Physician-Scientist Scholar Program to D.A.S.

